# Therapeutic targeting of Syndecan-1 axis overcomes acquired resistance to KRAS-targeted therapy in gastrointestinal cancers

**DOI:** 10.1101/2024.08.06.606865

**Authors:** Mitsunobu Takeda, Madelaine S. Theardy, Alexey Sorokin, Oluwadara Coker, Preeti Kanikarla, Shuaitong Chen, Zecheng Yang, Phuoc Nguyen, Yongkun Wei, Jun Yao, Xiaofei Wang, Liang Yan, Yanqing Jin, Yiming Cai, Masakatsu Paku, Ziheng Chen, Kara Z. Li, Francesca Citron, Hideo Tomihara, Sisi Gao, Angela K. Deem, Jun Zhao, Huamin Wang, Samir Hanash, Ronald A DePinho, Anirban Maitra, Giulio F. Draetta, Haoqiang Ying, Scott Kopetz, Wantong Yao

## Abstract

The therapeutic benefit of recently developed mutant KRAS (mKRAS) inhibitors has been limited by the rapid onset of resistance. Here, we aimed to delineate the mechanisms underlying acquired resistance to mKRAS inhibition and identify actionable targets for overcoming this clinical challenge. Previously, we identified Syndecan-1 (SDC1) as a key effector for pancreatic cancer progression whose surface expression is driven by mKRAS. By leveraging both pancreatic and colorectal cancer models, we found that surface SDC1 expression was initially diminished upon mKRAS inhibition, but recovered in tumor cells that bypass mKRAS dependency. Functional studies showed that these tumors depended on SDC1 for survival, further establishing SDC1 as a driver for the acquired resistance to mKRAS inhibition. Mechanistically, we revealed that the YAP1-SDC1 axis was the major driving force for bypassing mKRAS dependency to sustain nutrient salvage machinery and tumor maintenance. Specifically, YAP1 activation mediated the recovery of SDC1 localization on cell surface that sustained macropinocytosis and enhanced the activation of multiple RTKs, promoting resistance to KRAS-targeted therapy. Overall, our study has provided the rationale for targeting the YAP-SDC1 axis to overcome resistance to mKRAS inhibition, thereby revealing new therapeutic opportunities for improving the clinical outcome of patients with KRAS-mutated cancers.

## Introduction

KRAS is the most frequently mutated oncogene in human cancers^1^. Mutant KRAS (KRAS*) is critical for disease initiation in numerous cancer types, such as pancreatic ductal adenocarcinoma (PDAC) and non-small cell lung cancer (NSCLC), and accordingly, is detectable in early neoplastic lesions and remains functional in invasive and metastatic disease. Due to its high prevalence in cancer types and central role in tumor progression, the effective and specific targeting of KRAS* has been a continuing priority in anti-cancer drug development. Attempts to target KRAS* and other key components of the RAS-MAPK/RAS-PI3K pathway have long produced limited clinical response due to the feedback activation of compensation pathways or high drug toxicity. Notably, recent efforts have resulted in the development of highly selective KRAS^G12C^ inhibitors (KRAS^G12C^i), including Adagrasib (MRTX849)^2^ and Sotorasib (AMG510)^3^, which have been approved by the U. S. Food and Drug Administration (FDA) for the treatment of KRAS^G12C^-mutated NSCLC. However, the duration of therapy response remains short-lived due to the rapid development of acquired resistance. With agents targeting other KRAS mutants, such as KRAS^G12D^, entering clinical trials, there is an urgent need to improve our understanding of the molecular mechanisms driving acquired resistance to KRAS*-targeted therapies, as well as develop novel strategies for broadening the applicability of KRAS* inhibitors.

Recent studies using preclinical models have indicated that acquired resistance to KRAS^G12C^ inhibition may be mediated by the activation of multiple receptor tyrosine kinase (RTK) regulators, including EGFR, FGFR, IGF1R, HER2 and SHP2, as well as activation of RAS effectors, such as MYC and mTOR^2, 4–8^. Using a genetically engineered mouse (GEM) model of PDAC driven by inducible KRAS^G12D^, we have recently demonstrated that the activation of the YAP1 oncogene drives resistance to the genetic inactivation of KRAS*, which has been further confirmed in KRAS*-driven CRC^9–11^. However, the molecular mechanisms underlying the re-activation of the RTK-RAS signaling pathway as well as the contribution of YAP1 activation to developing resistance to KRAS* inhibitors remain to be elucidated. Clarification of the effector pathways contributing to KRAS* inhibitor resistance will provide opportunities for improving the efficacy and duration of KRAS*-targeted therapies.

Recently, we identified Syndecan-1 (SDC1), a cell surface proteoglycan, as a key effector downstream of KRAS* and demonstrated that KRAS*-driven SDC1 membrane expression plays a critical role in PDAC progression^12^. Here we aimed to gain a more complete understanding of the role of SDC1 in the regulation of KRAS*-dependency. We found that surface SDC1 expression was tightly correlated with acquired resistance to genetic or pharmacological inhibition of KRAS* in both PDAC and CRC preclinical models, and confirmed that the YAP1 oncogene is the major driver for SDC1 reactivation in cells resistant to KRAS* inhibition. Further, our findings elucidated a critical role for the YAP1-SDC1 axis for activating multiple RTKs to establish acquired resistance to KRAS* blockade. Our findings support the translational potential of targeting the YAP1-SDC1 axis to overcome the resistance to KRAS*-targeted therapies.

## Results

### SDC1 membrane expression is resumed upon acquired resistance to KRAS*-MAPK signaling blockade

Previously, we demonstrated that spontaneous tumor relapse following *Kras^G12D^* extinction, in doxycycline-inducible KRAS^G12D^-driven (iKras) GEM models of PDAC, was driven by KRAS*-independent mechanisms (KRAS*-Escapers; E–) or oncogene reactivation (KRAS*-reactivated; E+)^9^. Interestingly, while *Kras^G12D^* extinction resulted in a dramatic downregulation of cell surface SDC1^12^, membrane SDC1 level was comparable between E+ and E– tumors from the iKras model or tumor-derived primary cultures, although E– tumors exhibited much weaker MAPK activity than the E+ tumors **(Figure 1A-B)**. These data suggest that the recovery of membrane SDC1 in E– cells likely involved mechanisms beyond reactivation of KRAS*-MAPK signaling. The recovery of membrane SDC1 levels in these KRAS*-bypass cells prompted us to characterize the dynamic change of SDC1 membrane expression upon KRAS^G12D^ inactivation in the mouse iKras tumor cells. As we previously reported^12^, *Kras^G12D^* inactivation upon doxycycline withdrawal resulted in rapid and continuous downregulation of surface SDC1 levels during the first week. However, surface SDC1 expression gradually re-emerged following long-term KRAS* extinction **(Figure 1C, S1A)**. The bidirectional change of surface SDC1 expression following KRAS^G12D^ extinction conformed with the initial loss and eventual restoration of proliferative capacity, as indicated by Ki-67, of iKras tumor cells (**Figure S1B**).

**Figure 1.**
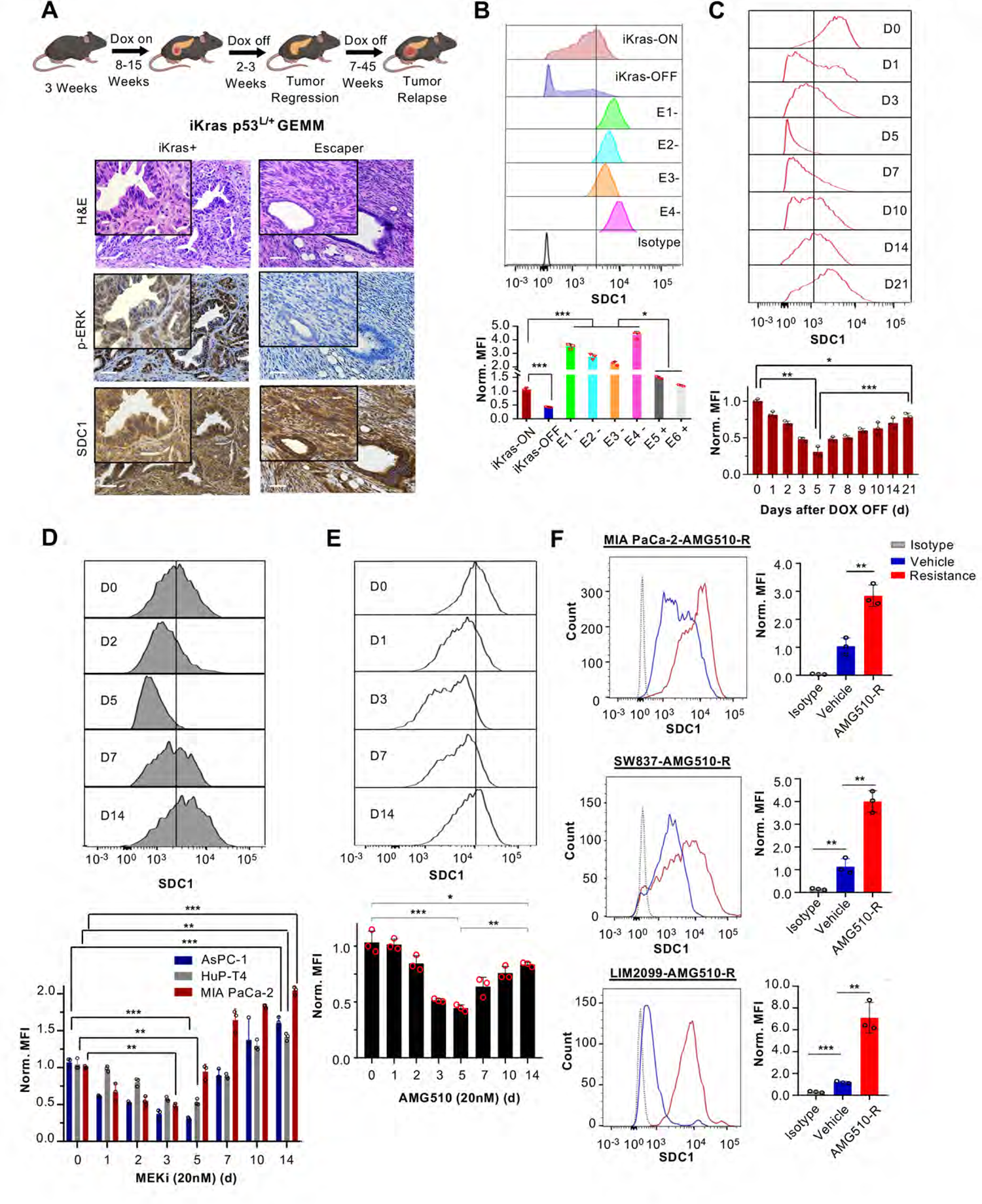
SDC1 membrane expression recovers upon acquired resistance to KRAS*-MAPK signaling blockade. (A) Experimental design for the development of PDAC tumors resistant to KRAS* blockade from iKras p53^L/+^ mouse models (top). H&E staining and immunohistochemistry for SDC1 and p-ERK of iKras as well as iKras-relapse escaper tumors (bottom). Scale bar, 100 µm. (B) SDC1 expression levels in iKras iKras p53^L/+^ tumor cells in the presence (ON) or absence (OFF) of doxycycline as well as in iKras-escaper lines (E1-E4). n = 3 biological replicates. (C) Surface SDC1 expression levels of iKras p53^L/+^ tumor cells (AK10965) upon doxycycline withdrawal for the indicated days. n = 3 biological replicates. (D) Human PDAC cells, AsPC-1, HuP-T4 and MIA PaCa-2, were treated with trametinib (20nM), for the indicated time periods, and surface protein levels of SDC1 were measured. n = 3 biological replicates. (E) Human PDAC MIA PaCa-2 cells cultured were treated with AMG510 (20nM) for the indicated time periods, and surface protein levels of SDC1 were measured. n = 3 biological replicates. (F) Surface SDC1 expression levels of AMG510-R cells derived from MIA PaCa-2, SW837, or LIM2099 cells chronically treated with AMG510 or vehicle control. n = 3 biological replicates. *p < 0.05, **p < 0.01, ***p < 0.001. Data are presented as mean ± SD.

To investigate the role of surface SDC1 in KRAS*-driven tumors under a clinically relevant context, we evaluated SDC1 surface expression upon pharmacological inhibition of KRAS* or its downstream MAPK pathway. Overall, we observed a bidirectional change in surface SDC1 expression consistent with SDC1 changes observed upon the genetic extinction of *Kras^G12D^* in mouse iKras tumor cells. Specifically, the initial depletion and subsequent restoration of surface SDC1 accumulation were observed upon inhibition of MAPK signaling using the MEK inhibitor, Trametinib, in both mouse iKras tumor cells and human PDAC cell lines, including MIA PaCa2, AsPC-1, and HuP-T4 cells (**Figure 1D**, **S1C**). This bidirectional change of surface SDC1 expression was also observed upon treatment with the KRAS^G12C^ inhibitor, AMG510, in KRAS^G12C^-mutant PDAC (MIA PaCa2) and CRC (SW837, LIM2099) cells (**Figure 1E**, **S1D**), but not in AMG510-treated non-responsive AsPC-1(KRAS^G12D^) or HuP-T4 (KRAS^G12V^) cells (**Figure S1E-F**). To further validate the induction of surface SDC1 expression during adaptation to KRAS* inhibition, we established AMG510-resistant (AMG510-R) cells derived from PDAC (MIA PaCa2) and CRC (SW837, LIM2099) cell lines that were chronically treated with AMG510 (**Figure S1G** and data not shown). Notably, surface SDC1 levels were significantly upregulated in AMG510-R cells compared to those parental cells (**Figure 1F, S1H** and data not shown), further strengthening the association between surface SDC1 expression and the emergence of KRAS* independence.

### SDC1 reconstitution enables tumor growth independent of the oncogenic KRAS-MAPK pathway

To assess the role of SDC1 in KRAS* bypass, we first tested the capacity of enforced SDC1 expression to enable KRAS*-independent tumor growth. Using two independent iKras tumor cell lines, iKras cells transduced with a green fluorescent protein expression vector (GFP; negative control) failed to generate tumors in the absence of doxycycline (no KRAS* transgene expression), whereas iKras cells transduced with either Kras^G12V^ (positive control) or SDC1 expression vectors generated tumors in the absence of doxycycline (**Figure 2A**). Notably, iKras cells expressing soluble SDC1 (SDC1 lacking cytoplasmic and transmembrane domains) were not able to establish tumor growth. This indicates that the localization of SDC1 to the membrane is critical to bypass KRAS* (**Figure 2A**). Correspondingly, SDC1 depletion with shRNA in primary cultures derived from the KRAS*-inactivated and SDC1-expressing iKras cells (SDC1-bypass [SB] cells; **Figure S2A-C**) abolished clonogenic activity (**Figure 2B, S2D**). Consistent phenotypes were attained by CRISPR-mediated deletion of SDC1 (**Figure S2E-F**). Similarly, genetic depletion of SDC1 abolished the clonogenic activity of E– tumor cells, as well as of E+ cells derived from iKras relapsed tumors **(Figure S2G)**, indicating that SDC1 expression is essential for the in vitro growth of both KRAS*-independent and KRAS*-dependent cancer cells. Together, these in vitro and in vivo data demonstrate that SDC1 expression is necessary and sufficient to drive cancer cell growth and to maintain tumorigenic potential in the absence of KRAS* transgene expression, findings consistent with a role for SDC1 in the development of KRAS* inhibitor resistance.

**Figure 2.**
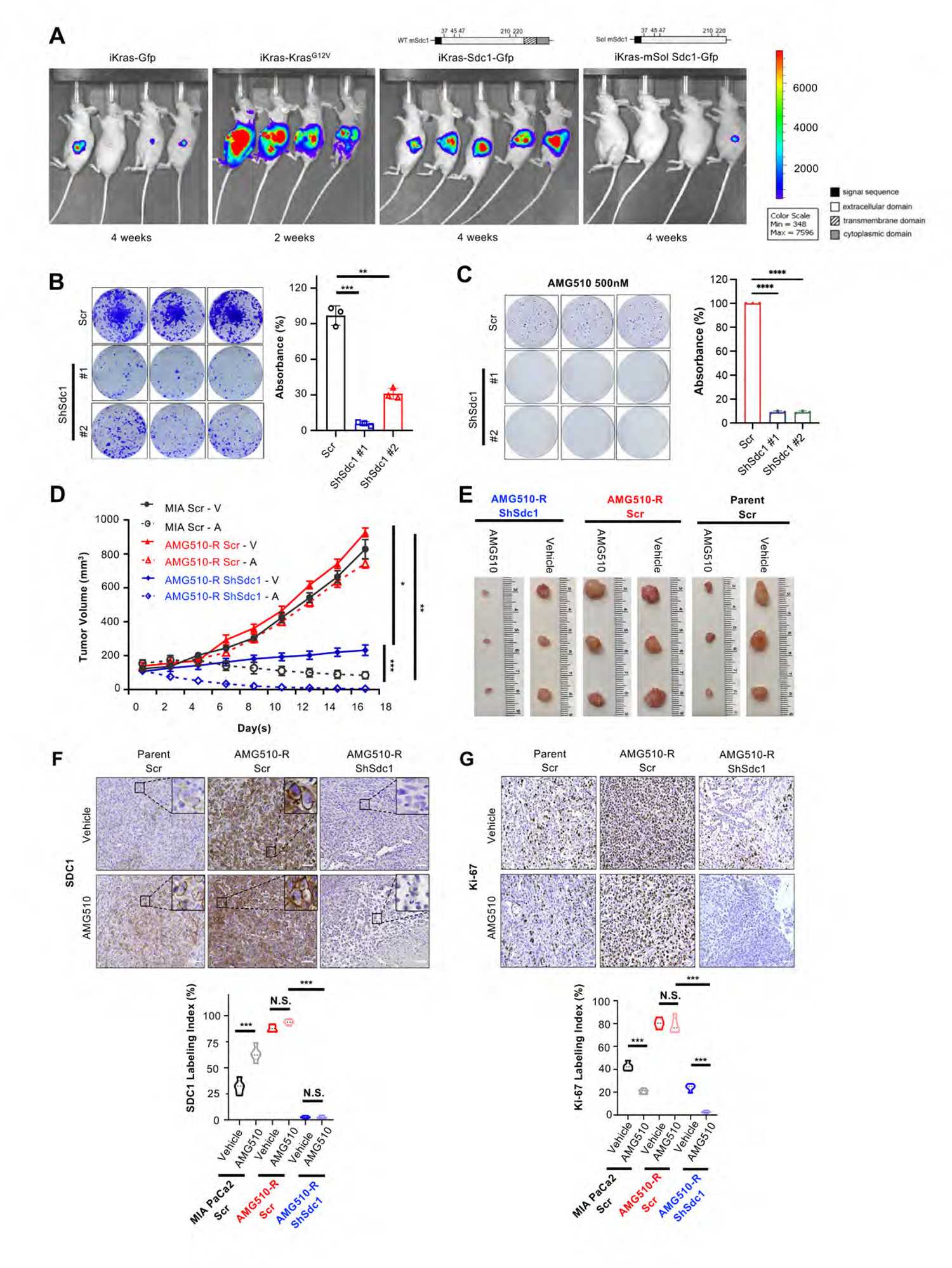
SDC1 reconstitution enables tumor growth independent of oncogenic KRAS-MAPK pathway. (A) Orthotopic xenografts generated from iKras cells. Post-implantation, mice were kept on DOX for 7 days and then switched to off DOX (to induced Kras^G12D^ inactivation). Tumor growth was visualized by bioluminescent imaging at 4 weeks off DOX for Gfp-, WT Sdc1-, and mSol Sdc1-expressing groups or at 2 weeks for the KRAS^G12V^-expressing group. (B) Clonogenic growth assay of SDC1 knockdown AK192 (192) cells derived from SDC1-bypass cells. n = 3 biological replicates. (C) Clonogenic growth assay of parental MIA PaCa-2 cells and their derived AMG510-R cells upon shRNA-mediated knockdown of Sdc1 in the presence of 500nM AMG510. n = 3 biological replicates. (D) Subcutaneous tumor growth of parental MIA PaCa-2 or AMG510-R cells transduced with shScramble or shSDC1, treated with AMG510 (A, 30 mg/kg) or Vehicle (V). Treatment started when tumors reached to ∼150mm^3^ and tumor sizes were measured every two days. n = 3 per group. (E) Representative photographs of dissected tumors from mice of Fig 2D. (F, G) Immunohistochemistry analysis for SDC1 (F) and Ki67 (G) in tumors from Fig 2E. Scale bar, 50 µm. Quantification was from ten random fields. *p < 0.05, **p < 0.01, ***p < 0.001, n.s. = non-significant. Data are presented as mean ± SD.

From these findings, we sought to assess whether SDC1 depletion could sensitize KRAS*-driven tumors to targeted therapy against KRAS* or MAPK signaling. SDC1 depletion with shRNA significantly sensitized mouse iKras tumor cells to trametinib *in vitro* as well as synergized with trametinib to significantly prolong survival of orthotopic xenograft models implanted with iKras tumor cells (**Figure S2H-J**). Similarly, SDC1 depletion sensitized human PDAC cells to trametinib or AMG510 *in vitro* (**Figure S2K-L**). Consistent with findings using E– cells derived from iKras relapsed tumors (**Figure S2D**), SDC1 depletion markedly impaired colony formation and viability of AMG510-resistant PDAC and CRC cells treated with AMG510 (**Figure 2C, S 2M-N**). In the subcutaneous xenograft model of MIA PaCa2 cells, the parental cells were highly sensitive to AMG510 treatment but AMG510-R cells, established in vitro, remained completely resistant in vivo (**Figure 2D-E**), with striking expression of SDC1 and high proliferation ability as indicated by Ki67 (**Figure 2F-G**). Importantly, SDC1 knockdown in combination with AMG510 treatment significantly blunted the growth of AMG510-R tumors and resulted in complete tumor regression (**Figure 2D-G**). Together, our findings support an integral role for SDC1 in enabling emergence of resistance to KRAS* genetic extinction or pharmacologic inhibition, providing rationale for development of SDC1-targeted therapies for tumors with acquired resistance to KRAS*-targeted therapy.

### SDC1 mediates the activation of multiple RTKs in KRAS*-bypass tumors

To gain mechanistic insights into SDC1-mediated bypass of KRAS*-dependency, Reverse Phase Protein Array (RPPA) profiling was conducted on primary cancer cell lines derived from SB tumors as well as their parental GFP- or SDC1-expressing iKras cancer cells cultured in the presence or absence of doxycycline for 24h. Unsupervised clustering indicated that protein expression levels in SB cells derived from two individual iKras cells were distinct with those from parental KRAS*-driven cells (**Figure 3A**). While MAPK signaling components, such as pERK and c-RAF, were downregulated in SB cells, these cells exhibited signatures of epithelial-mesenchymal transition (EMT), including epithelial markers E-Cadherin and β-catenin as well as induction of the mesenchymal markers Snail, N-cadherin, and collagen (**Figure 3A**). These patterns are reminiscent of the downregulation of MEK/MAPK signaling and induction of EMT in E– tumors^9^ (**Figure S3A**). Notably, many proteins that were highly overexpressed in SB cancer cells were involved in growth factor signaling, including multiple RTKs such as EGFR, PDGFRβ, c-MET, HER2, and IGF1R (**Figure 3A**). These RPPA findings were further confirmed by Phospho-RTK Array and Western blot analyses showing the hyperactivation of multiple RTKs as well as the relative downregulation of MAPK and activation of AKT signaling in SB cells compared to the parental iKras cell (**Figure 3B, S3B**). Similarly, hyperactivation of RTKs was observed in E– cells from the iKras model (**Figure S3C**) and AMG510-R cells (**Figure 3C**). Importantly, depletion of SDC1 in SB cells (**Figure S3D**) and AMG510-R cells (**Figure S3E**) significantly suppressed the activation of multiple RTKs as well as downstream AKT-mTOR signaling (**Figure 3C-E, S3F-H**). Together, these data indicate that SDC1 functions to sustain RTK signaling which is consistent with clinical observations of RTK activation in patients with acquired resistance to KRAS*-targeted therapy^8^.

**Figure 3.**
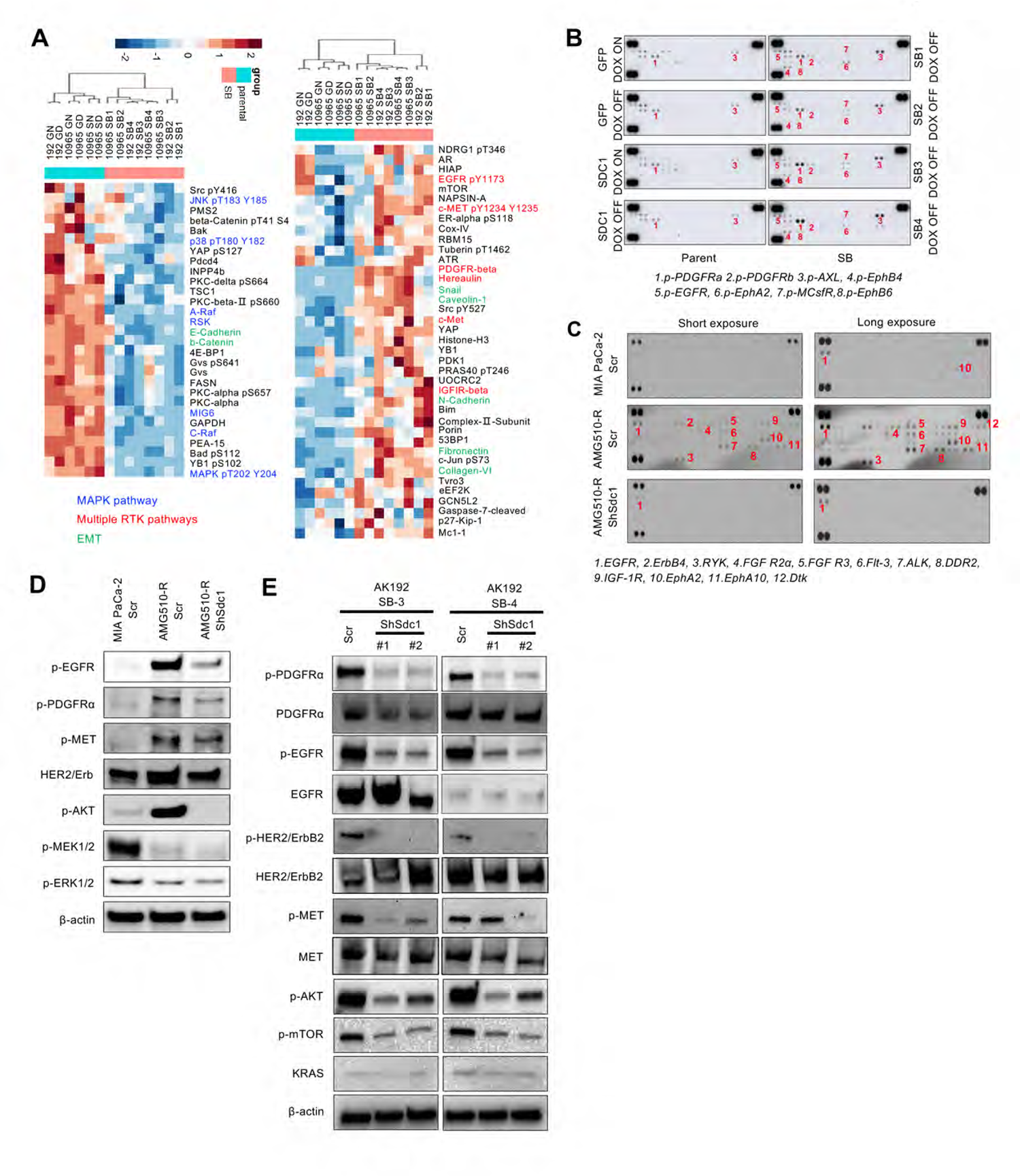
SDC1 mediates the activation of multiple receptor tyrosine kinases (RTKs) in KRAS-bypass tumors. (A) RPPA heatmap of genes enriched in SDC1-bypass (SB) cells compared to parental cells (GN (GFP-no DOX for 24 hr), GD (GFP- on DOX), SN (SDC1-no DOX for 24 hr), SD (SDC1-on DOX)). Expression levels shown are representative of log2 values of each replicate. (B) Phospho-RTK array analysis of parental and SB cells as described in Figure 3A. (C) Phospho-RTK array analysis of MIA PaCa-2 and AMG510-R infected with scrambled shRNA or shRNA against Sdc1. (D) Western blot analysis of MIA PaCa-2 and AMG510-R cells infected with scrambled shRNA or shRNA against Sdc1. (E) Western blot analysis of SB cells derived from individual mouse tumor (192 SB-3 and 192 SB-4) and infected with scrambled shRNA or shRNA against Sdc1.

### YAP1 drives the recovery of SDC1 expression through inhibition of Guanine-Nucleotide Activating Protein (GAP) to reactivate ARF6 activity

We previously demonstrated that the reduction of SDC1 membrane expression induced upon KRAS* inactivation was regulated through the control of SDC1 trafficking and independent of SDC1 transcription/expression^12^. Similarly, SDC1 protein levels remained unchanged during the recovery of membrane SDC1 levels in iKras cells that acquired resistance to KRAS* inactivation in vitro (**Figure S4A**). In contrast, the activity of ARF6, a small GTPase drives KRAS*-mediated SDC1 membrane recycling^12^, was initially decreased and eventually recovered in the iKras cells following long-term KRAS* inactivation or in E– cells (**Figure 4A-B, S4B**). This finding suggests that the recovery of surface SDC1 levels in KRAS*-bypass cells was likely mediated by trafficking mechanism. We previously demonstrated that ARF6 activity in KRAS*-driven PDAC cells was controlled by PSD4, an ARF6 guanine-nucleotide exchange factor (GEF) with transcription suppressed upon KRAS or MEK inactivation^12^. Interestingly, PSD4 expression remained low in KRAS*-bypass cells (**Figure 4C, S4C**), indicating that PSD4 was unlikely the driver of ARF6 activation and surface SDC1 recovery. To better understand the mechanisms mediating ARF6 reactivation during the acquisition of resistance to KRAS* inhibition, we mined the expression profiles of E– and E+ cells^9^ for ARF6-specific GEFs and GAPs. Our data show that the expression of several ARF6-specific GAPs, including ASAP2, ARAP2, ARAP3, was significantly decreased in E– compared to E+ cells (**Figure 4D**). This was further validated by Western blot analysis, which revealed that the downregulation of ASAP2 and ARAP2 closely mirrored the recovery of surface SDC1 during acquired resistance to KRAS* inhibition in mouse iKras cells and human MIA PaCa-2 cells (**Figure 4C&E**). The downregulation of GAPs upon acquired resistance to the inhibition of KRAS* signaling was also observed in human PDAC cells following long-term MEK inhibition (**Figure S4C**). Importantly, reconstitution of ASAP2 expression in AMG510-R cells indeed diminished surface SDC1 levels (**Figure 4F-G**), underscoring the critical role of ASAP2 downregulation for SDC1 recovery during acquired resistance to KRAS inhibition.

**Figure 4.**
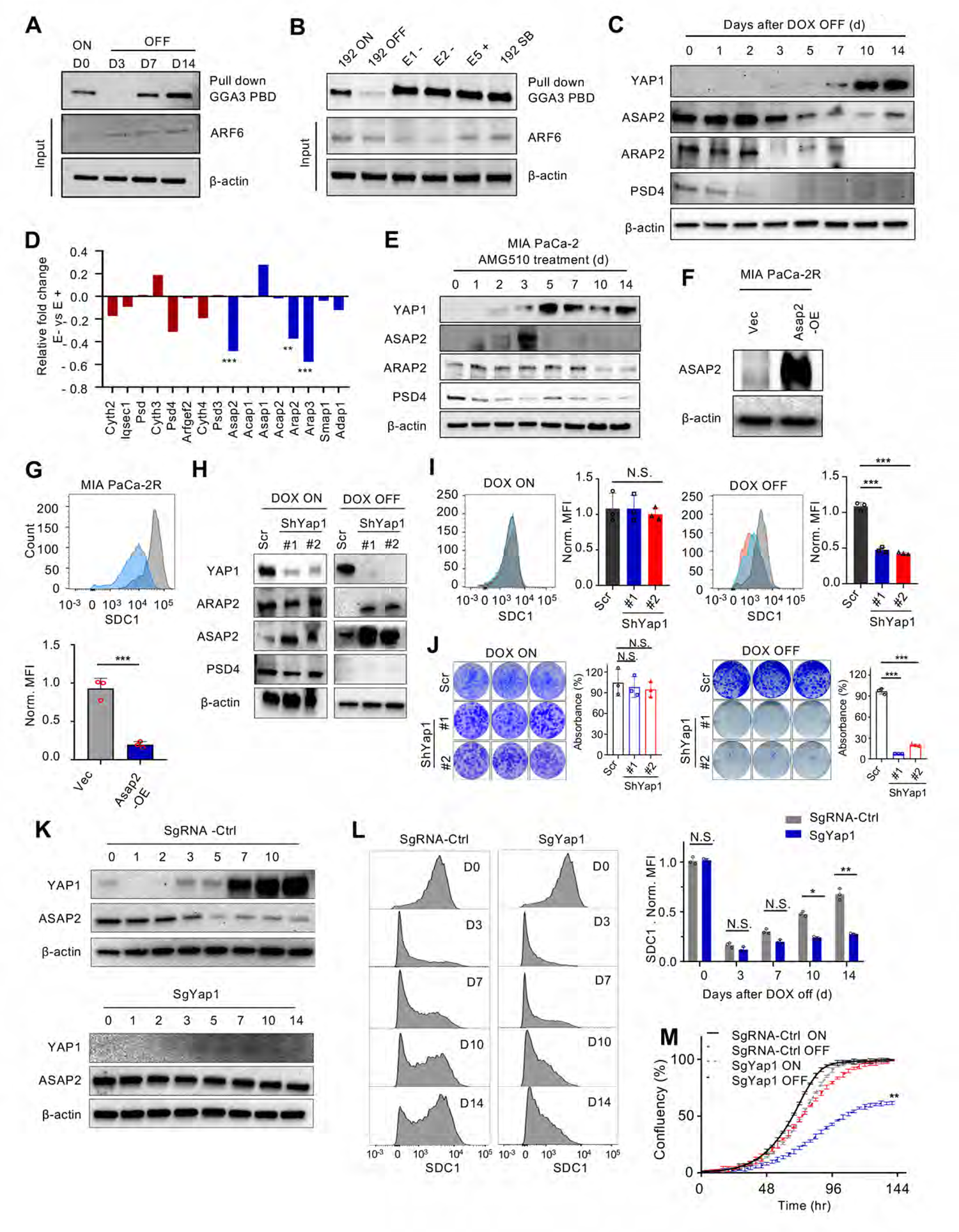
YAP1 drives the recovery of SDC1 expression through repressing guanine-nucleotide activating protein (GAP) to reactivate ARF6 activity. (A, B) ARF6 activity was measured in iKras p53^L/+^ tumor cells (192) grown in the presence (ON) or absence (OFF) of doxycycline (DOX) for indicated days (A), as well as E+, E-, and SB cells. Input lysates were immunoblotted to validate expression of ARF6 and β-actin. (C) Western blot analysis of iKras p53^L/+^ tumor cells (AK192) grown in the absence of DOX for indicated time periods. (D) mRNA expression of ARF6 GAPs and GEFs in the E+ and E-cell microarray dataset during KRAS-dependent and -independent relapse. n = 4 biological replicates. (E) Western blot analysis of MIA PaCa-2 cells treated with AMG510 (20 nM) for indicated time periods. (F) Cell lysates of MIA PaCa-2 AMG510-R cells expressing ectopic Asap2 (Asap-OE) or empty vector (Vec) were immunoblotted for ASAP2 and β-actin. (G) Surface SDC1 expression levels of MIA PaCa-2R expressing ectopic Asap2 (Blue peak) or empty vector (grey peak) were measured. n = 3 biological replicates. (H) Western blot analysis of iKras p53^L/+^ tumor cells (AK192, DOX ON and OFF) infected with scrambled shRNA or shRNA against Yap1. (I) Surface SDC1 expression levels of iKras p53^L/+^ tumor cells (DOX ON and OFF) infected with scrambled shRNA (grey peak) or shRNAs against Yap1 (blue and red peaks). n = 3 biological replicates. (J) Clonogenic growth assay of iKras p53^L/+^ tumor cells (AK192) infected with scrambled shRNA or shRNA against Yap1 upon KRAS* activation (DOX ON) or KRAS* inactivation (DOX OFF for 21 days). n = 3 biological replicates. (K) Western blot analysis of iKras p53^L/+^ tumor cells with CRISPR/Cas9-mediated Yap1 knockout (sgYAP1) or with non-targeting sgRNA (sgRNA-Ctrl) grown in absence of DOX for indicated time periods. (L) Surface SDC1 expression levels of iKras p53^L/+^ tumor cells (Ctrl) or their derived cells with CRISPR/Cas9-mediated Yap1 knockout (sgYAP1) grown in absence of DOX for indicated days. n = 3 biological replicates. (M) iKras p53^L/+^ tumor cells with Cas9-mediated Yap1 knockout (sgYAP1) or sgRNA control (sgRNA-Ctrl) were grown in the presence (ON) or absence (OFF) of DOX. Cell proliferation was measured using the live-cell time-lapse imaging module of Incucyte. n = 6 biological replicates. *p < 0.05, **p < 0.01, ***p < 0.001, n.s. = non-significant. Data are presented as mean ± SD.

We further sought to elucidate the molecular mechanisms underlying ASAP2 downregulation during acquired resistance to KRAS inhibition. We had identified YAP1 activation, through either gene amplification or activation of a non-canonical WNT pathway, as the major driver for bypassing KRAS dependency in PDAC ^9, 11^. Interestingly, in the iKras PDAC mouse cells or MIA PaCa-2 cells following long-term genetic or pharmacological inactivation of KRAS*, or human PDAC cells upon long-term MEK inhibition, accumulation of YAP1 protein occurred in parallel with a decrease in ASAP2 and ARAP2 expression (**Figure 4C&E, S4C)**. While YAP1 knockdown did not affect ARAP2 or ASAP2 levels in the presence of KRAS*, nor did clonogenic activity of KRAS* dependent cells, depletion of YAP1 dramatically induced the expression of both ARAP2 and ASAP2, depleted surface SDC1 levels as well as suppressed clonogenic activity of the KRAS* independent cells (**Figure 4H-J, S4D-E**). These findings were further confirmed by using iKras cells with CRISPR-Cas9-mediated *Yap1* knockout, which demonstrated that *Yap1* deletion abolished downregulation of ARAP2 or ASAP2 expression (**Figure 4K**) and blocked the recovery of surface SDC1 levels (**Figure 4L**) following KRAS* inhibition. Importantly, YAP1 ablation significantly diminished the proliferative and clonogenic activity of KRAS* independent, but not parental KRAS*-dependent cells (**Figure 4M, S4F**). These data suggest that YAP1 directly or indirectly mediates the downregulation of ARF6 GAPs to promote SDC1 expression on the plasma membrane during acquired resistance to KRAS* inhibition.

### YAP1-SDC1 is required for macropinocytic activity of cells resistant to KRAS* inhibition

The ability to stimulate macropinocytosis, a regulated form of endocytosis, is a distinctive feature of KRAS* activation^13^, and PDAC cells harboring KRAS* rely on increased levels of macropinocytosis for nutrient salvaging to sustain uncontrolled cell growth^14^. Our previous study demonstrated that the surface localization of SDC1 driven by KRAS* activation played a crucial role in maintaining macropinocytosis in PDAC^12^. To better understand the function of SDC1 and the requirement of micropinocytosis for acquiring resistance to KRAS* inhibitors, we examined the macropinocytic activity in KRAS* bypass tumors. Although extinction of KRAS* leads to the rapid reduction of SDC1 surface expression and macropinocytosis^12^, macropinocytosis levels in E– tumor cells were similar to that in E+ cells and iKras cells (**Figure 5A-B**), suggesting that macropinocytosis is recovered in those escapers through a KRAS*-independent mechanism. The high macropinocytic level in the context of KRAS* bypass was confirmed in MIA PaCa2 AMG510-R resistant cells. MIA Paca2 parental and AMG510-R cells were grown in low-glutamine conditions and addition of albumin can rescue cell viability, while macropinocytosis inhibition by EIPA treatment reverses this effect indicating that macropinocytosis-mediated albumin uptake is critical for the viability of both KRAS*-dependent and -independent cells (**Figure 5C**). Accordingly, EIPA treatment diminished colony forming in AMG510-R cells, underscoring the essential role of macropinocytosis for proliferation in the context of KRAS*-resistance (**Figure 5D**).

**Figure 5.**
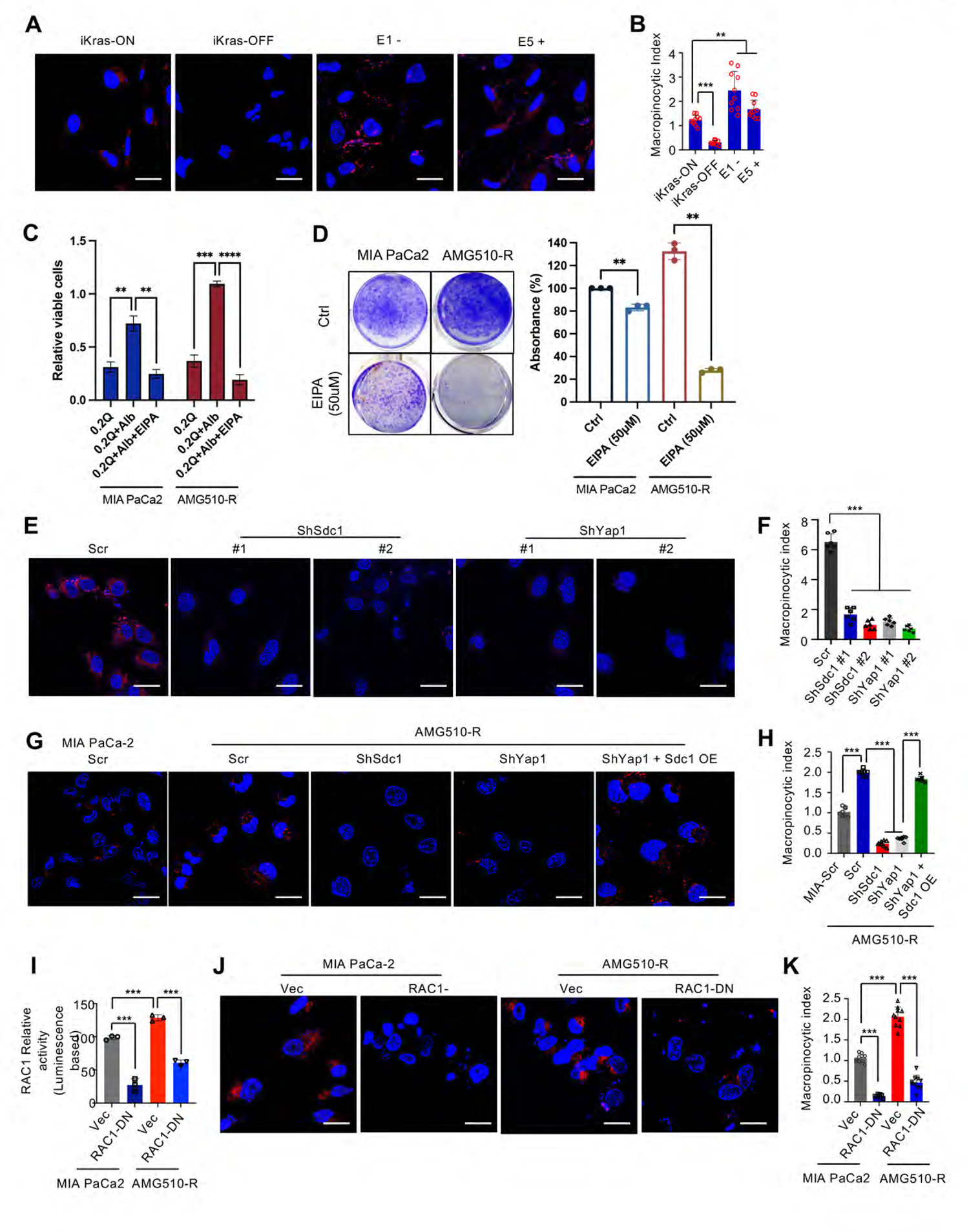
The YAP1-SDC1 axis is required for macropinocytic activity in tumor cells resistant to KRAS* inhibition. (A, B) Macropinocytosis was visualized (A; scale bar, 20 µm) and quantified (B; n = 10 areas) in iKras p53^L/+^ tumor cells as well as E– and E+ cells grown in the presence (ON) or absence (OFF) of doxycycline for 3 days. (C) Glutamine deprivation assay of MIA PaCa2 parental and AMG510-R cells. Cells are grown in either one of the following conditions: 0.2Q (0.2 mM glutamine), 0.2mM glutamine with 2% albumin (Alb), or 0.2mM glutamine and 2% Alb with 25 µM EIPA treatment. Values are presented as relative number of viable cells at the time endpoint of the assay. (D) Clonogenic growth formation assay of parental and AMG510-R MIA PaCa2 cells treated with either DMSO control or 50 µM EIPA. (E, F) Macropinocytosis was visualized (E; scale bar, 20 µm) and quantified (F; n = 10 areas) in E– cells infected with two different Sdc1 and Yap1-targeting shRNA or scrambled shRNA control. (G, H) MIA PaCa-2 and AMG510-R cells stably expressing ectopic Sdc1 or empty vector were infected with scrambled shRNA or shRNA against Sdc1 or Yap1. Macropinocytosis was visualized (G; scale bar, 20 µm) and quantified (H; n = 10 areas). (I) RAC1 activity in MIA PaCa-2 and AMG510-R cells stably expressing Rac1-dominant negative (RAC1-DN) or empty vector (Vec). n = 3 biological replicates. (J, K) Macropinocytosis in MIA PaCa-2 and AMG510-R cells stably expressing Rac1-dominant negative (RAC1-DN) or empty vector (Vec) was visualized (J; scale bar, 20 µm) and quantified (K; n = 10 areas). *p<0.05, **p < 0.01, ***p < 0.001, ****p < 0.0001, n.s. = non-significant. Data are presented as mean ± SD.

Our data suggests that macropinocytosis is recovered in the E– cells possibly through KRAS*-independent but SDC1-dependent mechanisms. Indeed, ablation of SDC1 abolished macropinocytic activity in E– cells and AMG510-resistant MIA PaCa-2 cells (**Figure 5E-H, S 5A-B**), indicating that SDC1 was required for macropinocytosis in these KRAS*-bypass cells. Interestingly, YAP1 knockdown also decreased surface SDC1 and suppressed macropinocytic activity in these KRAS*-escaper cells (**Figure 5E-F, S5A-B**), which could be rescued by ectopic expression of SDC1 (**Figure 5G-H, S5B**). These findings, coupled with the observation that YAP1 mediates the downregulation of ARF6-GAPs to promote SDC1 surface localization, suggest that YAP1 may drive SDC1-dependent macropinocytosis in cells resistant to KRAS* inhibition. The small GTPase Rac1 plays a crucial role in the formation of initial membrane ruffles and macropinocytosis^15^. Our previous study has shown that RAC1 activity is mediated by SDC1 to promote macropinocytosis^12^. Here, we found that RAC1 activity is higher in E– and AMG510-R cells compared to that in iKras cells and parental MIA PaCa-2 cells, respectively (**Figure 5I, S5C**). In AMG510-R cells, ectopic expression of dominant negative RAC1 (RAC1-DN, RAC1(T27N)) suppressed macropinocytic activity but did not affect SDC1 expression (**Figure 5J-K, S5D-E**). Together, these data indicate that the SDC1-Rac1 axis is required for macropinocytosis in tumor cells resistant to KRAS*-targeted therapy.

### Targeting the YAP1-SDC1 axis abolishes acquired resistance to KRAS* inhibition

To evaluate the translational potential of targeting the YAP1-SDC1 axis, we employed therapeutic antisense oligonucleotides (ASO(s), IONIS Pharmaceuticals) that target YAP1. YAP1-ASO treatment depleted YAP1 protein levels in both parental MIA PaCa-2 and SW837 cells as well as the corresponding AMG510-R cells (**Figure 6A, S6A, B, K**). However, YAP1 knockdown only induced ASAP2 and ARAP2 expression (**Figure 6A, S6A, B, K**) and depleted surface SDC1 expression (**Figure 6B, S6C, D, L**) in a dose-dependent manner in AMG510-R cells. This is consistent with our findings from KRAS* bypass (**Figure 4H-I**) and AMG510-R cells (**Figure S6E-F**) transfected with YAP1-targeting shRNA. Moreover, YAP1 depletion with ASO treatment (**Figure 6C-D, S6M**) or shRNA (**Figure S6G-H**) abolished clonal formation activity in AMG510-R, but not their parental cells. These data suggest that the YAP1-SDC1 axis is specifically required for the in vitro growth of cancer cells with acquired resistance to KRAS* inhibition. Accordingly, depletion of YAP1 with YAP1-ASO in the subcutaneous xenograft model of MIA PaCa-2 cells resulted in decreased membrane SDC1 and tumor regression only in tumors derived from AMG510-R, but not parental, cells (**Figure 6E, S6I-J**). Moreover, YAP1-ASO treatment in parental MIA PaCa-2 cells blocked the restoration of surface SDC1 following KRAS* inhibition and significantly sensitized the cells to AMG510 treatment (**Figure S6N-O**), underscoring the potential of targeting YAP1 to prevent the acquired resistance to KRAS* inhibition.

**Figure 6.**
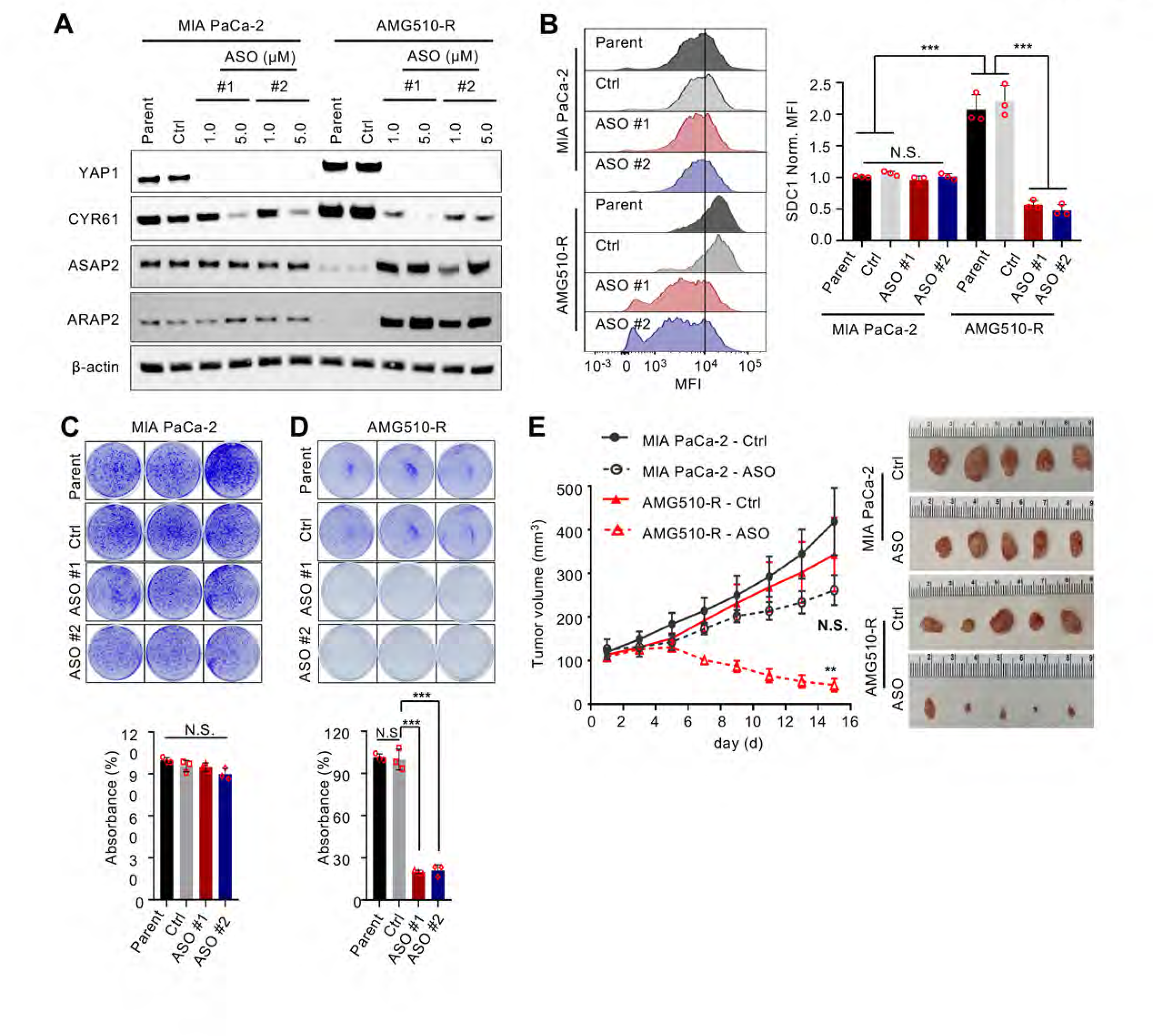
Targeting the YAP1-SDC1 axis abolishes acquired resistance to mutant KRAS-targeted therapy. (A) Western blot analysis of MIA PaCa-2 or AMG510-R cells treated with 1µM or 5µM of YAP1 ASOs or Control-ASO for 72 h. (B) Surface SDC1 expression levels of MIA PaCa-2 and AMG510-R cells treated with 5µM of YAP1-ASOs or Control-ASO. n = 3 biological replicates. (C, D) Clonogenic growth assay of MIA PaCa-2 (C) and AMG510-R (D) cells treated with control ASO (Ctrl) or YAP1-ASOs. (E) MIA PaCa-2 and AMG510-R cells were subcutaneously injected into 5-week-old female NGS mice. Mice were treated with Control ASOs (50 mg/kg, SQ, QD (5 days ON, 2 days OFF)) or YAP1-ASOs (50 mg/kg, SQ, QD (5 days ON, 2 days OFF)) when tumors size reached to 100 mm^3^. Tumor volumes were measured every two days (left) and representative photographs of dissected tumors are shown (right). Results are presented as the means ± SD (n=5). **p < 0.01, ***p < 0.001, * n.s. = non-significant. Data are presented as mean ± SD.

To assess the applicability of our findings in clinically relevant models, we evaluated responses to AMG510 in a cohort of CRC patient derived xenograft (PDX) models harboring KRAS^G12C^. The stratification of tumors sensitive (B8026, C1047) or moderately sensitive (C1177, F3008) to AMG510 treatment was conducted by monitoring the rate of tumor growth under AMG510 treatment (**Figure S7A**). To analyze the expression and function of the YAP1-SDC1 axis during acquired resistance to Kras^G12C^ inhibition, models with acquired resistance were developed from AMG510-sensitive CRC PDX tumors following long-term AMG510 treatment (B8026-R, C1177-R, F3008-R) (**Figure S7B-D**). While 70-90% of PDAC cases exhibit high SDC1 expression ^12^, parental CRC PDXs only exhibit low SDC1 expression (**Figure S7G**). In contrast, both SDC1 and YAP1 expression are highly induced in PDX tumors with acquired resistance to AMG510 (**Figure 7A, S7E-F**), thus underscoring the activation of YAP1-SDC1 axis during acquired resistance to KRAS*-targeted therapy. More importantly, YAP1-ASO treatment improved the sensitivity to AMG510 in all resistant PDX models, leading to significant tumor growth cessation or regression without overt toxicity (**Figure 7B, S7H-I**). Moreover, combined treatment with YAP1-ASO and AMG510 resulted in the ablation of SDC1 expression, MAPK and AKT signaling as well as YAP1 expression in AMG510-resistant PDX models, accompanied by the decrease in cell proliferation as indicated by Ki67 positivity and elevated apoptosis as indicated by cleaved caspase-3 (**Figure 7C-D, S7J-K**). This was further confirmed by RPPA analysis, which revealed diminished MAPK and YAP1 signaling upon corresponding treatment as well as the downregulation of multiple RTKs upon the combination treatment in tumors from each group collected at day 2 (**Figure 7E, S7L**) and day 21 (**Figure 7G, S7M**) after treatment. Moreover, consistent with our previous findings on the role of glycolysis as a critical downstream effector of oncogenic KRAS signaling in advanced tumors^16, 17^, combined treatment with YAP1-ASO and AMG510 led to a significant decrease of multiple glycolysis enzymes, including hexokinase (HK1, HK2), enolase (ENO1, ENO2), lactate dehydrogenase (LDH) and lactate transporter (MCT4), in samples collected at day 21 after treatment (**Figure 7F, S7M**). Together, our findings provide strong preclinical evidence for positioning YAP1-SDC1 blockade as a novel strategy for overcoming resistance to KRAS*-targeted therapy.

**Figure 7.**
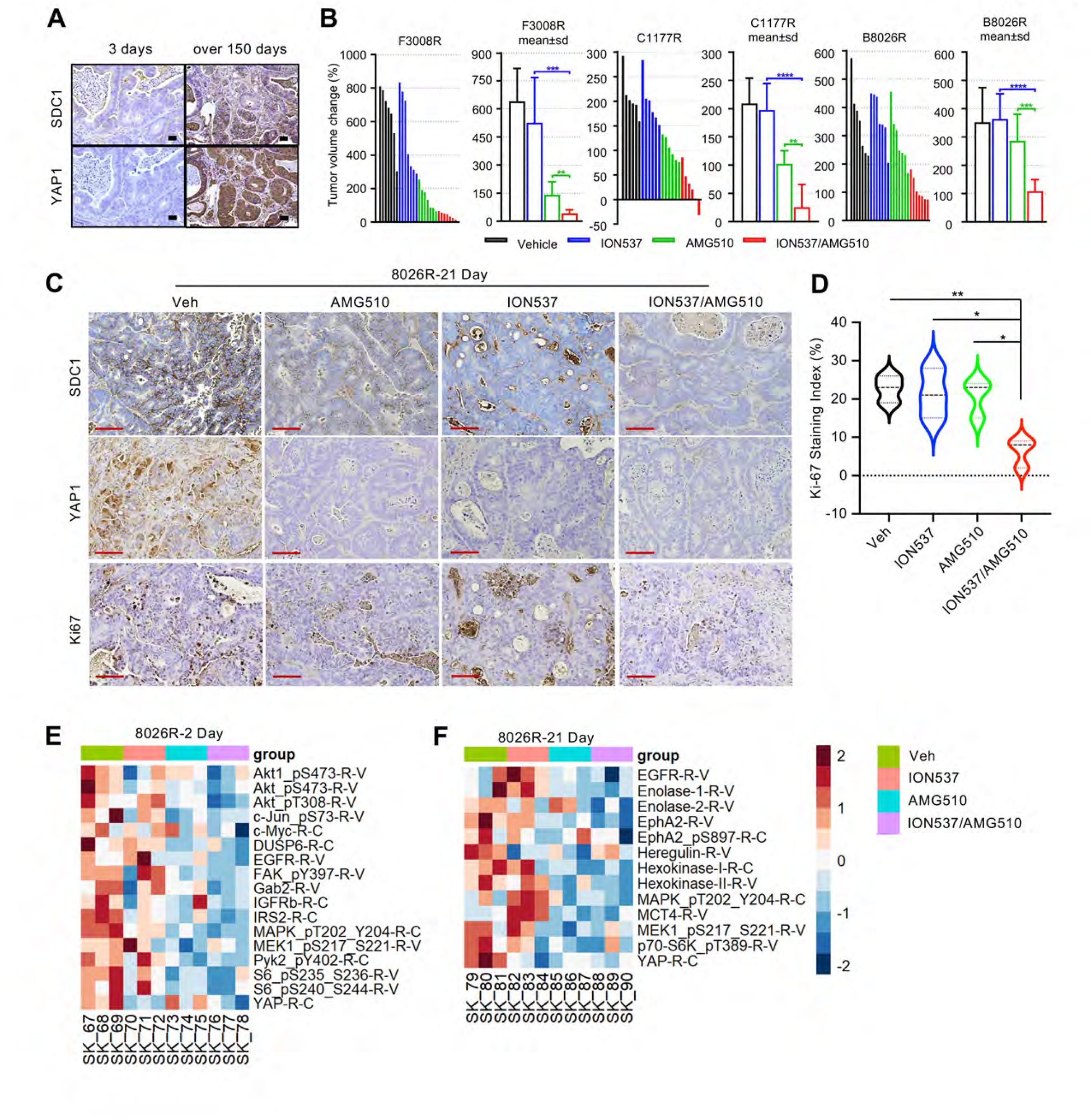
Targeting YAP1 reverses acquired resistance to mutant KRAS-targeted therapy in CRC PDX models. (A) Immunohistochemistry analysis of SDC1 and YAP1 expression in colorectal tumor tissue collected from models from Fig S7B at 3 days or over 150 days after clear progression (tumor volume change >100% from baseline) and after initiation of AMG510 treatment. Scale bar, 50 µm. (B) PDX models with acquired resistance to AMG510 were treated with Vehicle (Veh), YAP1-ASO (ION537; 50 mg/kg, SQ, QD (5 days ON, 2 days OFF)), AMG510 (30 mg/kg, PO, QD), or ION537 and AMG510 combination treatment. n = 10 per group. (C) Immunohistochemistry for SDC1, YAP1, and Ki67 expression in tumor tissue collected from models described in Fig 7B at 21 days after initiation of treatment. Scale bar, 100 µm. (D) Quantification of Ki-67 staining from Figure 7C. n = 3 per group. (E, F) RPPA heatmap of B8026R tumors, from at 2 days (E) or 21 days (F) of treatment. Protein levels shown are log2 values of each sample. *p<0.05, **p < 0.01, ***p < 0.001, ****p < 0.0001, n.s. = non-significant. Data are presented as mean ± SD.

## Discussion

While the recent development of small molecule inhibitors specific for oncogenic KRAS has revolutionized the treatment landscape for KRAS*-driven cancers, the efficacy and applicability of these therapies have been limited by the rapid onset of acquired resistance. For instance, AMG510 treatment in patients with *KRAS^G12C^*-driven NSCLC achieved a response rate of over 30%^18, 19^, but all patients developed therapeutic resistance that limited long-term survival^19^. Similarly, our previous work using an inducible *Kras^G12D^*-driven PDAC model demonstrated spontaneous tumor relapse in the majority of animals following genetic extinction of *Kras^G12D^* ^9^. Such acquired resistance to KRAS^G12D^ inhibition is expected to be also inevitable for the ongoing clinical trials of multiple KRAS targeted therapies. The vertical reactivation of the RAS pathway elements has been characterized in the majority of samples collected from patients who developed resistance to KRAS^G12C^ inhibitors^4, 8, 20, 21^. One major route leading to such recovery of RAS activity is the activation of RTKs through either genetic (e.g. gene amplification or fusion) or non-genetic mechanisms ^2, 4–8, 22–25^. Our study also revealed the activation of multiple RTKs in AMG510-R tumors upon characterization of AMG510-R tumors from mouse iKras-escaper mice and the CRC PDX models, which was consistent with a recent study using human cancer cell lines^6^. This finding, thus, suggests that targeting multiple RTKs rather than a single RTK is likely required to overcome acquired resistance to KRAS* inhibition. Previously, inhibiting multiple RTKs through adaptors like SOS and SHP2, which function downstream of RTKs to promote RAS activation, has shown promising results to attenuate acquired resistance to KRAS inhibition in preclinical studies^7, 22, 26^, but concerns about the therapeutic window and potential toxicity limit this approach. Further, early studies have indicated that the activation of FOXO transcription factors or downregulation of MYC can induce feedback activation of RTK transcription following the inhibition of PI3K/AKT or MEK inhibition^27, 28^, but these adaptive mechanisms are difficult to therapeutically target. Here, we focus on SDC1, a transmembrane proteoglycan, consisting of a membrane-embedded core protein and covalently linked glycosaminoglycan (GAG) chains, which are mostly composed of heparan sulfate (HS) and chondroitin sulfate (CS). The HS chain of SDC1 has been shown to be a co-receptor for growth factors and promote the activation of several RTKs such as EGFR, c-MET, FGFR, and IGF1R^29–32^. Accordingly, our data indicate that SDC1 functions as a common upstream regulator for the activation of multiple RTKs in tumor cells with acquired resistance to the inhibition of oncogenic KRAS. Additionally, SDC1 is localized to the cell surface and is dispensable for normal tissue homeostasis^33^. Together, these findings suggest that SDC1 may serve as an actionable target to suppress the activation of multiple RTKs and overcome resistance to KRAS*-targeted therapy.

Previously, we demonstrated that SDC1plays a critical role for oncogenic KRAS-mediated macropinocytosis^12^, a nutrient salvage process that supports oncogenic growth^14^. Here we showed that, while macropinocytosis and surface SDC1 levels rapidly decreased upon inhibition of oncogenic KRAS, SDC1-mediated macropinocytosis recovered in tumor cells upon acquired resistance to KRAS inhibition. The importance of macropinocytosis for gaining independence from oncogenic KRAS-driven tumor growth has been confirmed by a recent study demonstrating the necessity of macropinocytosis, induced by USP21-MARK3-mediated regulation of microtubule dynamics, for Kras^G12D^-independent tumor growth in an iKras PDAC model^34^. Macropinocytosis has also been shown to be important for developing resistance to tyrosine kinase inhibitors and standard care agents, such as gemcitabine and 5-FU^35, 36^. Together, these data suggest that SDC1-mediated macropinocytosis may function as a general mechanism underlying therapy resistance.

Localization of SDC1 to the plasma membrane is critical for its function in both RTK activation and macropinocytosis. Our previous study indicates that ARF6 activity downstream of oncogenic KRAS signaling controls SDC1 surface localization in PDAC cells^12^. To build upon this discovery, our current study has demonstrated that, during acquired resistance to KRAS inhibition, plasma membrane localization of SDC1 is driven by YAP1-mediated recovery of ARF6 activity. Further studies need to be conducted to further unravel mechanisms by which YAP1 transcriptionally regulates ARF6 GAPs. The YAP1 oncogene has been well-established as one of the major drivers of bypassing oncogene addiction and promoting acquired resistance to targeted therapies in several cancer types, including pancreatic cancer, lung cancer, colorectal cancer, melanoma, and glioblastoma^9, 10, 37–39^. Here, we show that, while YAP1 is not directly involved in the KRAS signaling cascade, it indirectly functions to sustain KRAS downstream signaling during acquired resistance to KRAS* inhibition by maintaining plasma membrane localization of SDC1 for the activation of multiple RTKs and induction of macropinocytosis. Although YAP1 is essential for embryonic development^40^, genetic studies in various mouse models suggest that YAP1 is dispensable for the homeostasis of normal cells in adults^41^. Together, these data underscore the potential of therapeutically targeting YAP1 as an innovative strategy for overcoming the acquired resistance to KRAS* inhibition. Indeed, depletion of YAP1 with YAP1-ASOs sensitized the CRC PDX models to AMG510 treatment without exerting overt toxicity in animals. Because the YAP1-ASOs in our study are currently under phase 1 trial in advanced solid tumors (NCT04659096) and multiple YAP1 inhibitors are under clinical development, we are confident that our findings have the potential to be quickly translated into the clinic for combination trials with iExosomes^42^ (NCT03608631) or small molecular inhibitors targeting oncogenic KRAS.

## Acknowledgements.

We thank Yohei Yoshihama for technical help, Ningping Feng and Tim Heffernan for providing instructions for in vivo treatment, Dave Aten for help with the graphical abstract. We would also like to thank our colleagues at the Institute for Applied Cancer Science (IACS), the Flow Cytometry and Cellular Imaging Core, the Sequencing and Microarray Facility, the Department of Veterinary Medicine, Medical Graphics & Photography at The University of Texas MD Anderson Cancer Center (Cancer Center Support Grant, CA016672). We thank all members of H.Y.’s, S.K.’s and G.F.D.’s lab members for discussion and reagents.

## Materials and Methods

### Cell culture

Established human cell lines were obtained from the American Type Culture Collection (ATCC), Leibniz Institute DSMZ, or Expasy, and cultured according to manufacturer’s protocols. Establishment of primary PDAC lines was performed as previously described ^9^. Additional details for cell culture are described in Supplementary Methods.

### In vivo studies

Allograft and xenograft studies were carried out in NSG mice or NCr Nude mice (Taconic) and were approved by the MD Anderson IACUC under protocol number 00001549 and 00001077. Additional details for animal studies are described in Supplementary Methods.

### Detection and quantification of macropinocytosis

The macropinocytic index was determined as previously described ^12^. Additional details are described in Supplementary Methods.

### Treatment of Patient derived xenograft models

All *in vivo* experiments utilizing PDXs were performed according to NCI NIH recommendations summarized in SOP50102. Additional details for animal studies are described in Supplementary Methods.

### Grant support

The research was supported by the Pancreatic Cancer Action Network-AACR Pathway to Leadership Grant (16-70-25-YAO), Pancreatic Cancer Action Network-Translational Research Grant (19-65-YAO), W81XWH-20-1-0598 (Department of Defense, DOD), 1R01CA272744 (NCI), RSG-22-017-01-CCB (ACS), generous philanthropic donations to the MD Anderson Cancer Center Dallas Living Legend Fund and Moon Shots Program, and University Cancer Foundation via the Institutional Research Grant program at the University of Texas MD Anderson Cancer Center to W.Y.; The Uehara Memorial Foundation postdoctoral fellowship, The Shinya Foundation for International Exchange of Osaka University Graduate School of Medicine Grant and The Cell Science Research Foundation Grant to M.T.; 1R01CA214793 (NCI) and HT94252311082 (DOD) to H.Y.; 5P01CA11796 (NCI) to H.W., H.Y., A.M., and R.A.D.; GI SPORE Grant (P50CA221707, NCI) to H.Y., A.M., and S.K.; The Flow Cytometry Core and the Department of Veterinary Medicine at the MD Anderson Cancer Center (MDACC) were supported by the Cancer Center Support Grant, CA016672. U54CA224065 (NCI), R01CA262805 (NCI), R01CA184843 (NCI), Stand Up 2 Cancer Catalyst Team Grant, generous philanthropic contributions to The University of Texas MD Anderson Cancer Center Moon Shots Program™ as well as The Del and Dennis McCarthy Distinguished Professorship to S.K. MDACC RPPA Core was supported by NCI Grant # CA016672 and Dr. Yiling Lu’s NIH R50 Grant # R50CA221675: Functional Proteomics by Reverse Phase Protein Array in Cancer. Sheikh Khalifa bin Zayed Foundation, and R01CA220236 (NCI) to A.M.

## Abbreviations

AMG510-R: AMG510-resistant
ASO: antisense oligonucleotide
CRC: colorectal cancer
CS: chrondroitin sulfate
EMT: epithelial-mesenchymal transition
GAG: glycosaminoglycan
GAP: GTPase-activating protein
GEF: guanine-nucleotide exchange factor
GEM: genetically engineered mouse
GFP: green fluorescence protein
HS: heparan sulfate
KRAS*: mutant KRAS
NSCLC: non-small cell lung cancer
ORR: overall response rate
PDAC: pancreatic ductal adenocarcinoma
PDX: patient-derived xenograft
RAC1-DN: dominant negative RAC1
RPPA: reverse phase protein array
RTK: receptor tyrosine kinase
SB: SDC1-bypass

## Disclosures

Dr. Scott Kopetz: Sanofi, Biocartis, Guardant Health, Array BioPharma, Genentech/Roche, EMD Serono, MedImmune, Novartis, Amgen, Lilly, Daiichi Sankyo. He serves as consultant for Roche, Genentech, EMD Serono, Merck, Karyopharm Therapeutics, Amal Therapeutics, Navire Pharma, Symphogen, Holy Stone, Biocartis, Amgen, Novartis, Lilly, Boehringer Ingelheim, Boston Biomedical, AstraZeneca/MedImmune, Bayer Health, Pierre Fabre, Redx Pharma, Ipsen, Daiichi Sankyo, Natera, HalioDx, Lutris, Jacobio, Pfizer, Repare Therapeutics, Inivata. Dr. Anirban Maitra: Dr. Maitra receives royalties from Cosmos Wisdom Biotech for a license related to a pancreatic cancer early detection test. He is also listed as an inventor on a patent licensed to Thrive Earlier Detection Ltd (an Exact Sciences Company) and serves as a consultant for Freenome and Tezcat Biotechnology. Dr. Ronald A. Depinho is currently Founder and Advisor of Tvardi Therapeutics, Asylia Therapeutics, Bectas Therapeutics, and Sporos Bioventures..

## Author Contributions

**Takeda M, Theardy MS**: Conceptualization, formal analysis, validation, investigation, visualization, methodology, writing–original draft, writing–review and editing. **Sorokin A, Coker O**: formal analysis, validation, investigation, visualization, methodology, writing–original draft, writing–review and editing. **Kanikarla P, Chen S, Yang Z, Nguyen P, Wei Y, Wang X, Yan L, Jin Y, Paku M, Chen Z, Li KZ, Citron F, Tomihara H, Zhao J**: Formal analysis, validation and investigation. **J. Yao**: Computational analysis, **Cai Y, Gao S, Deem AK**: visualization and writing–original draft. Formal analysis, validation and investigation. **Wang H**: Supervision and resources. **Hanash S, DePinho RA, Maitra A, Draeta GF**: Supervision, funding support and resources. **Ying H, Kopetz S, Yao W**: Conceptualization, resources, formal analysis, supervision, funding acquisition, validation, investigation, visualization, methodology, writing–original draft, project administration, writing–review and editing.

## Data transparency

All data is available upon request.

---

**Supplementary Figure 1.**
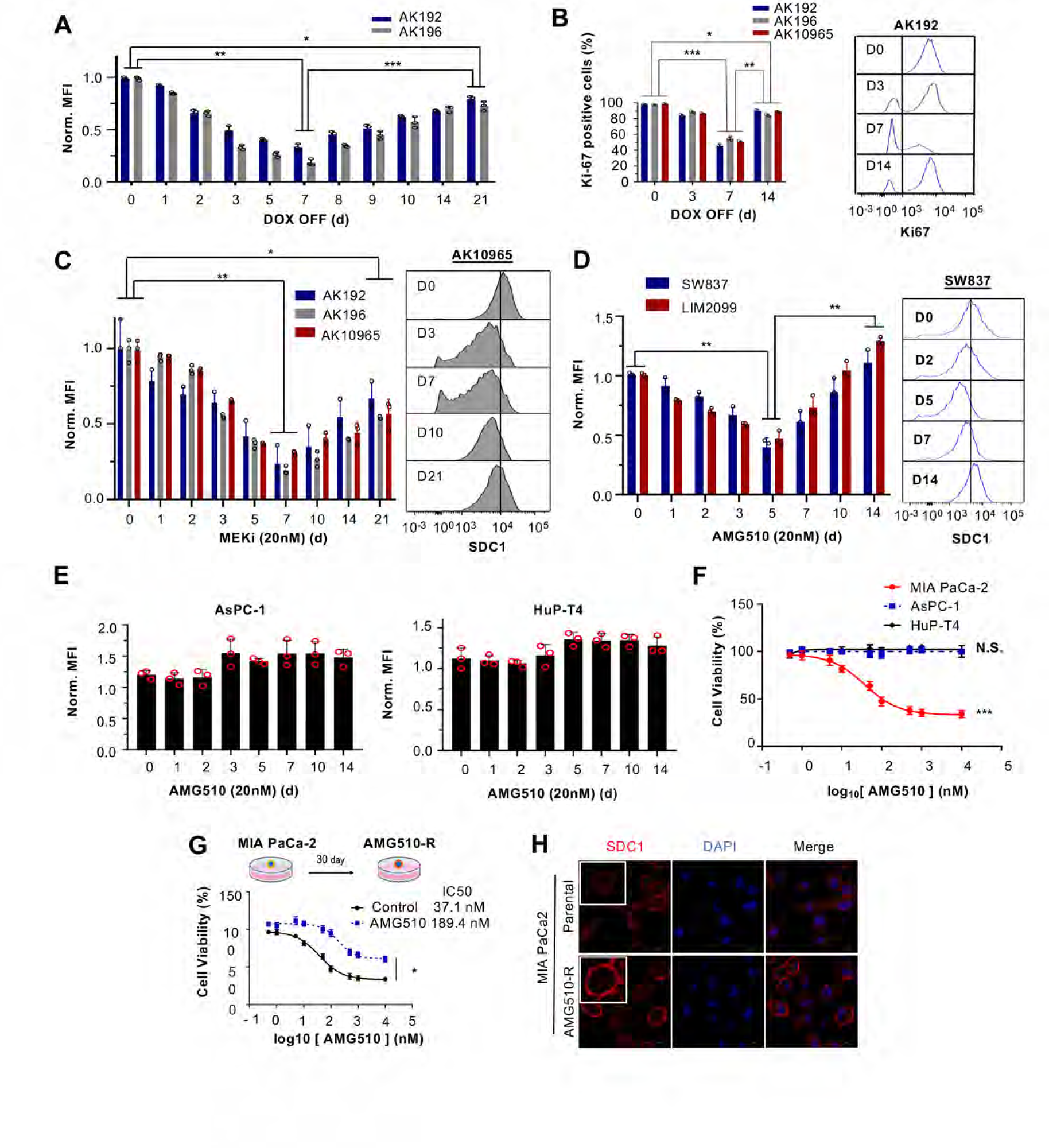

**Supplementary Figure 2.**
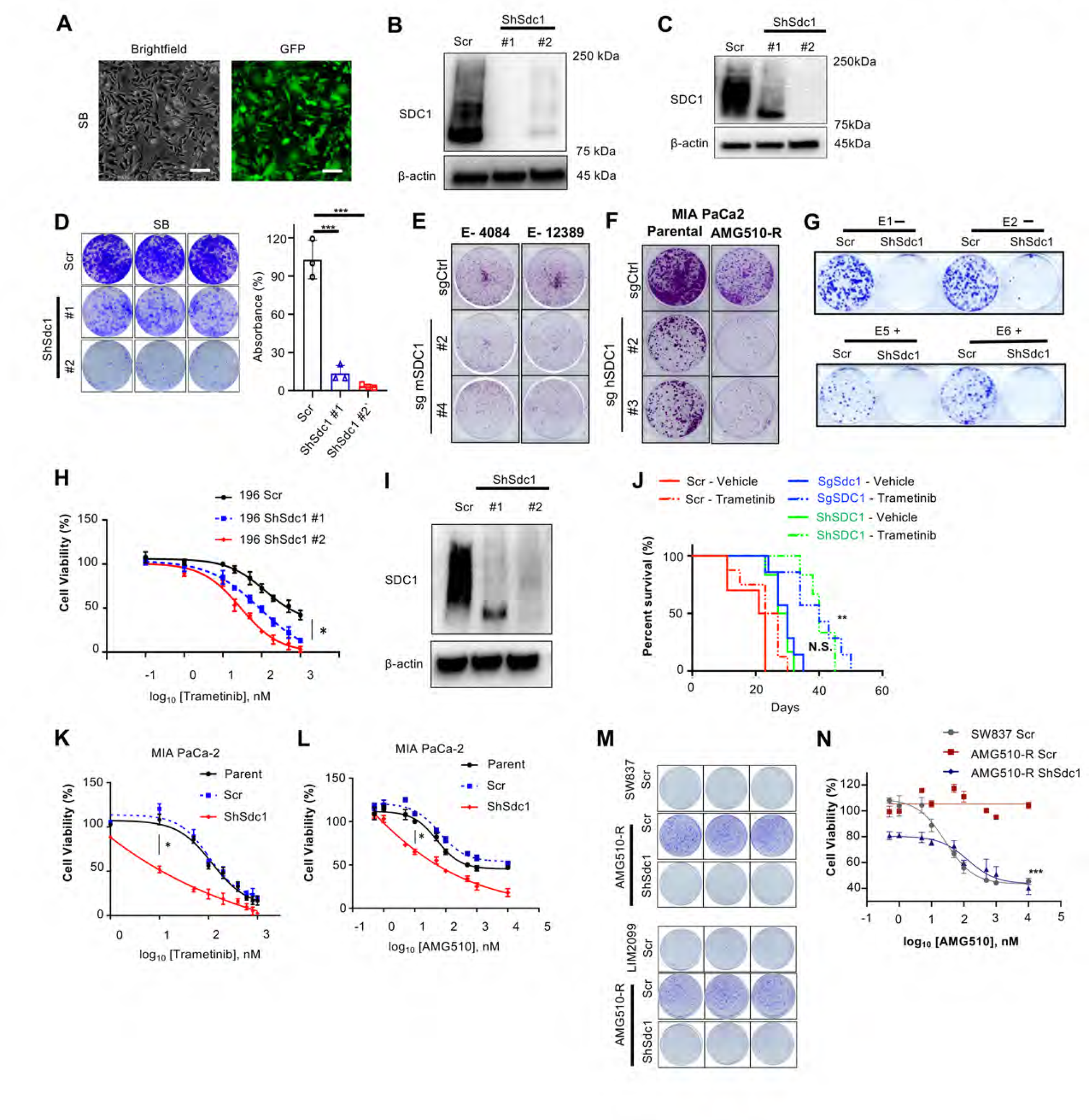

**Supplementary Figure 3.**
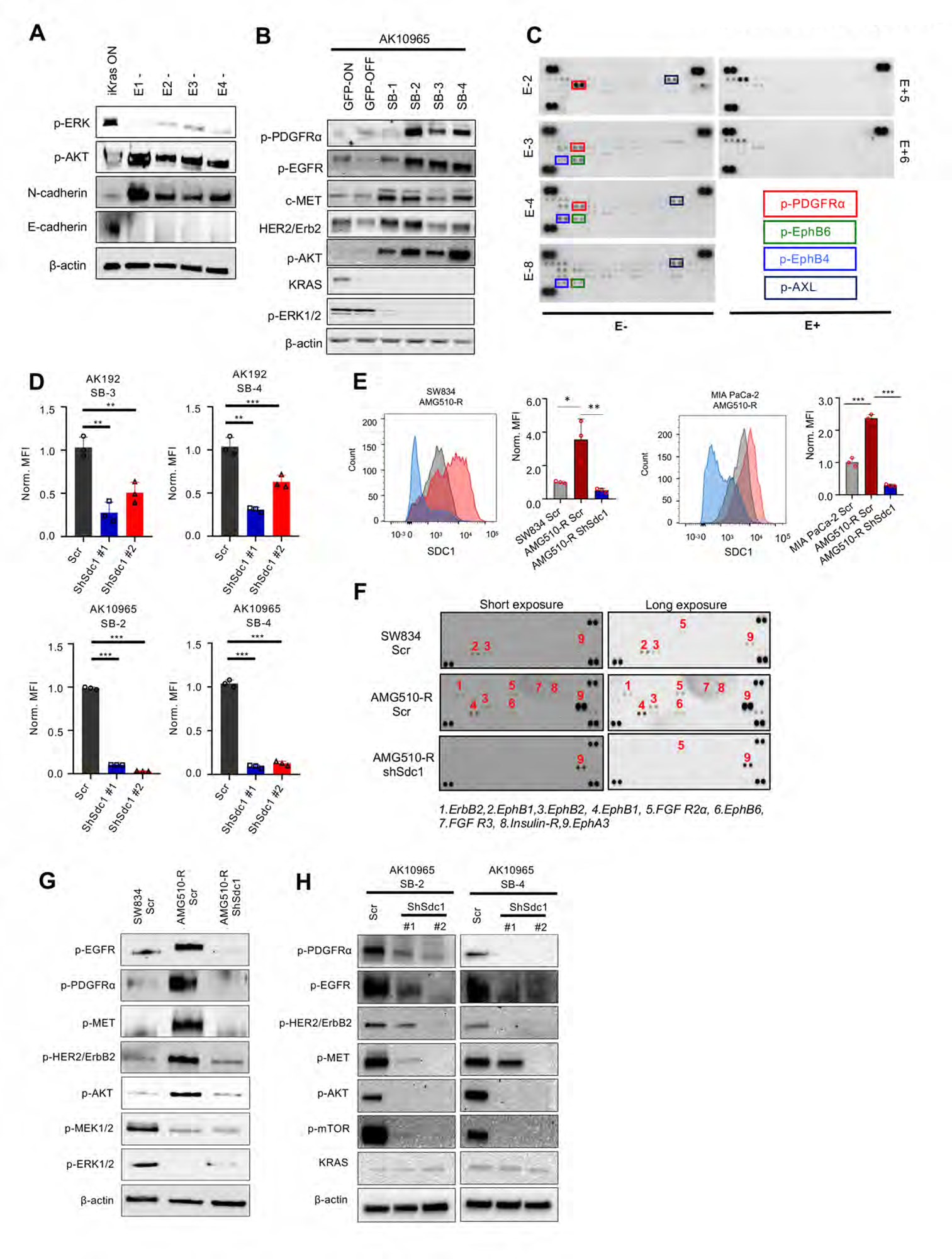

**Supplementary Figure 4.**
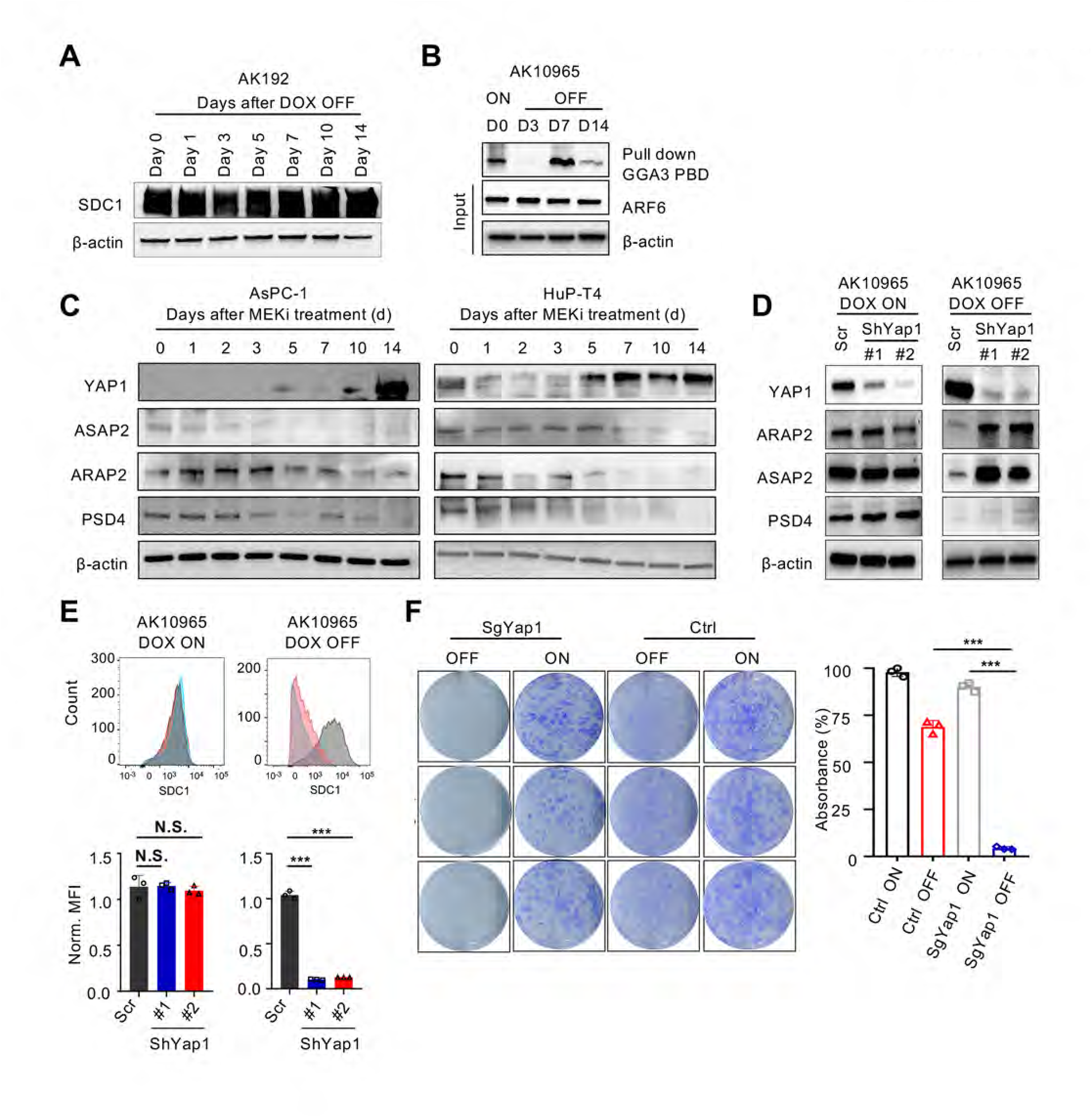

**Supplementary Figure 5.**
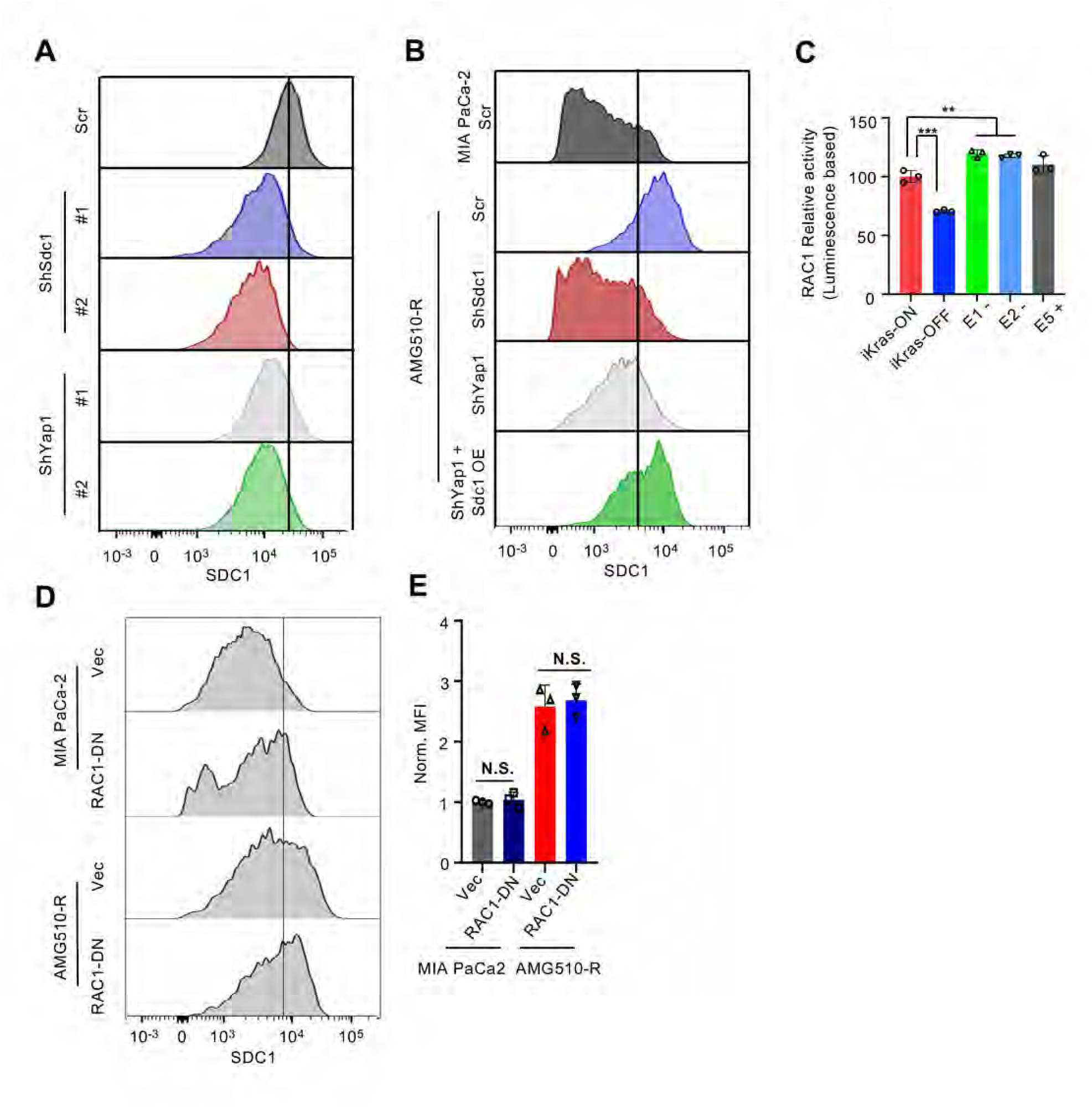

**Supplementary Figure 6.**
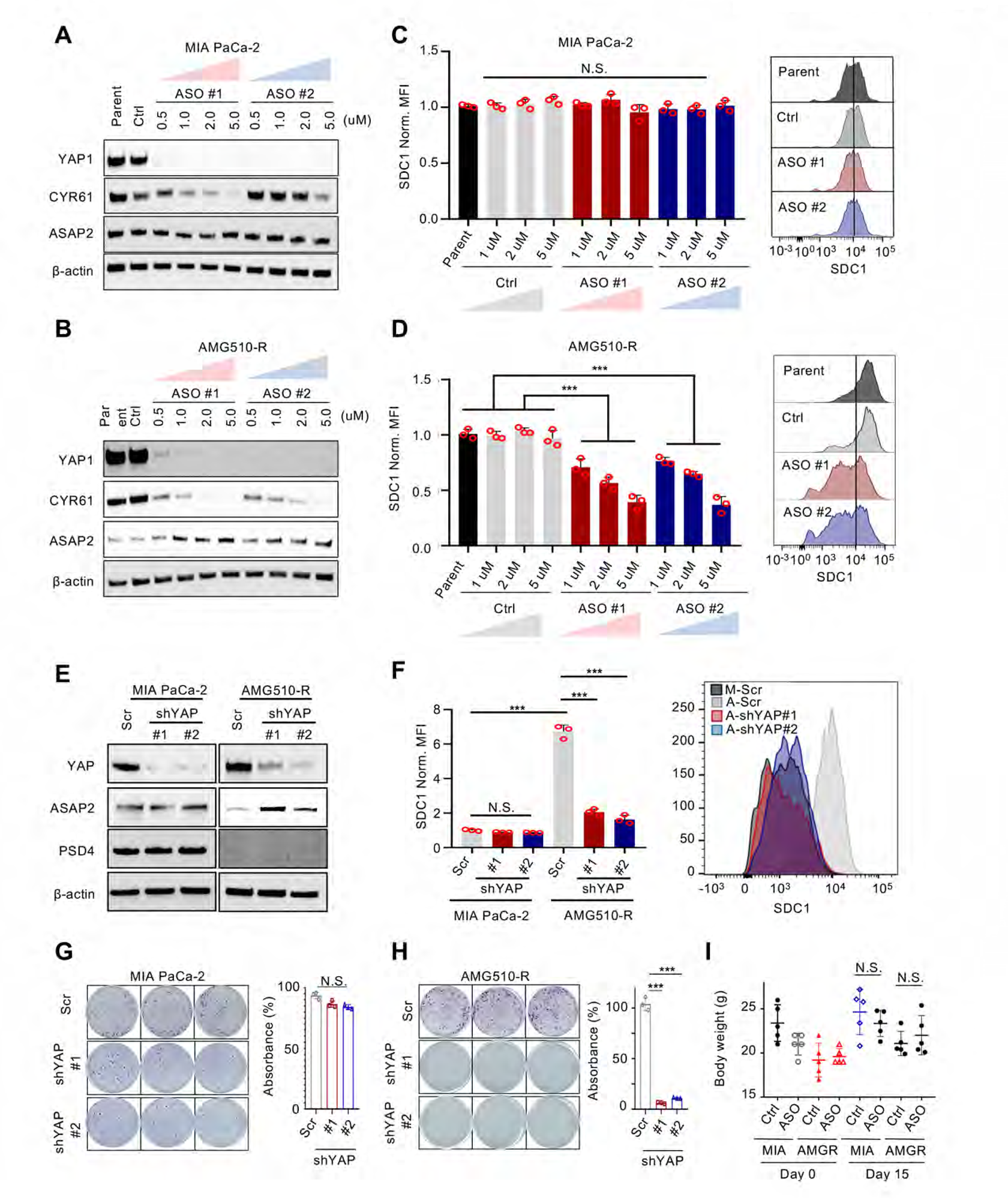
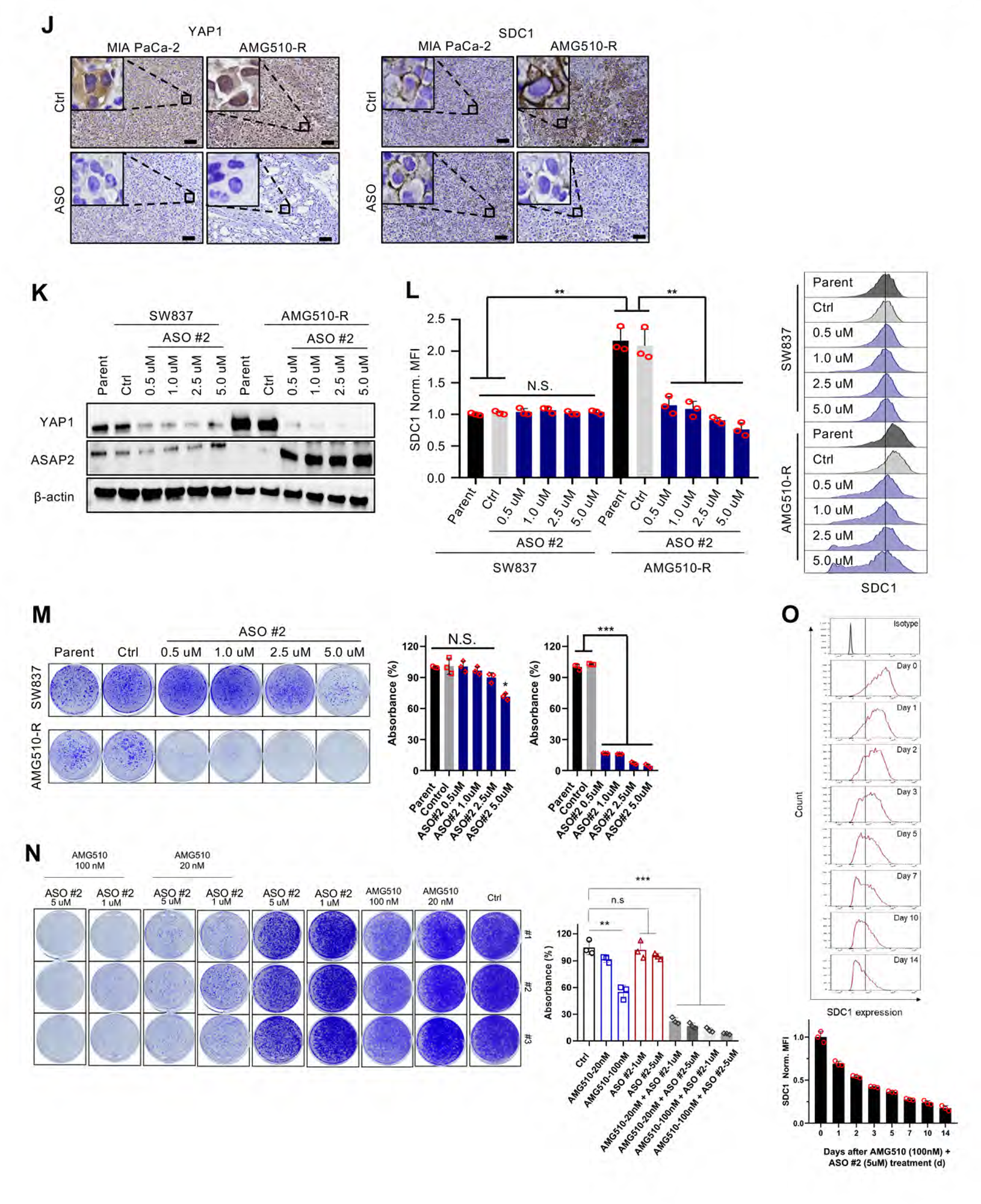

**Supplementary Figure 7.**
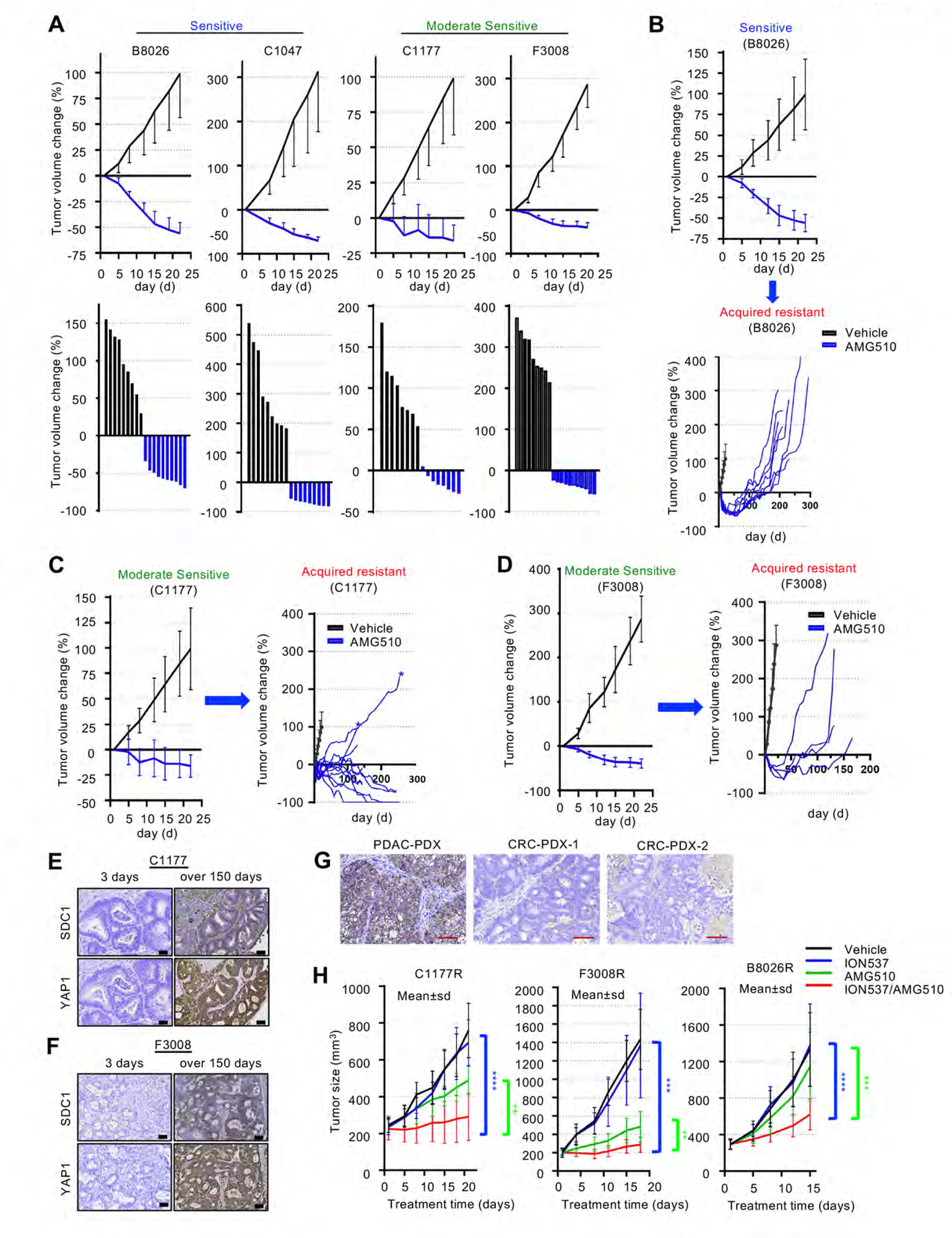
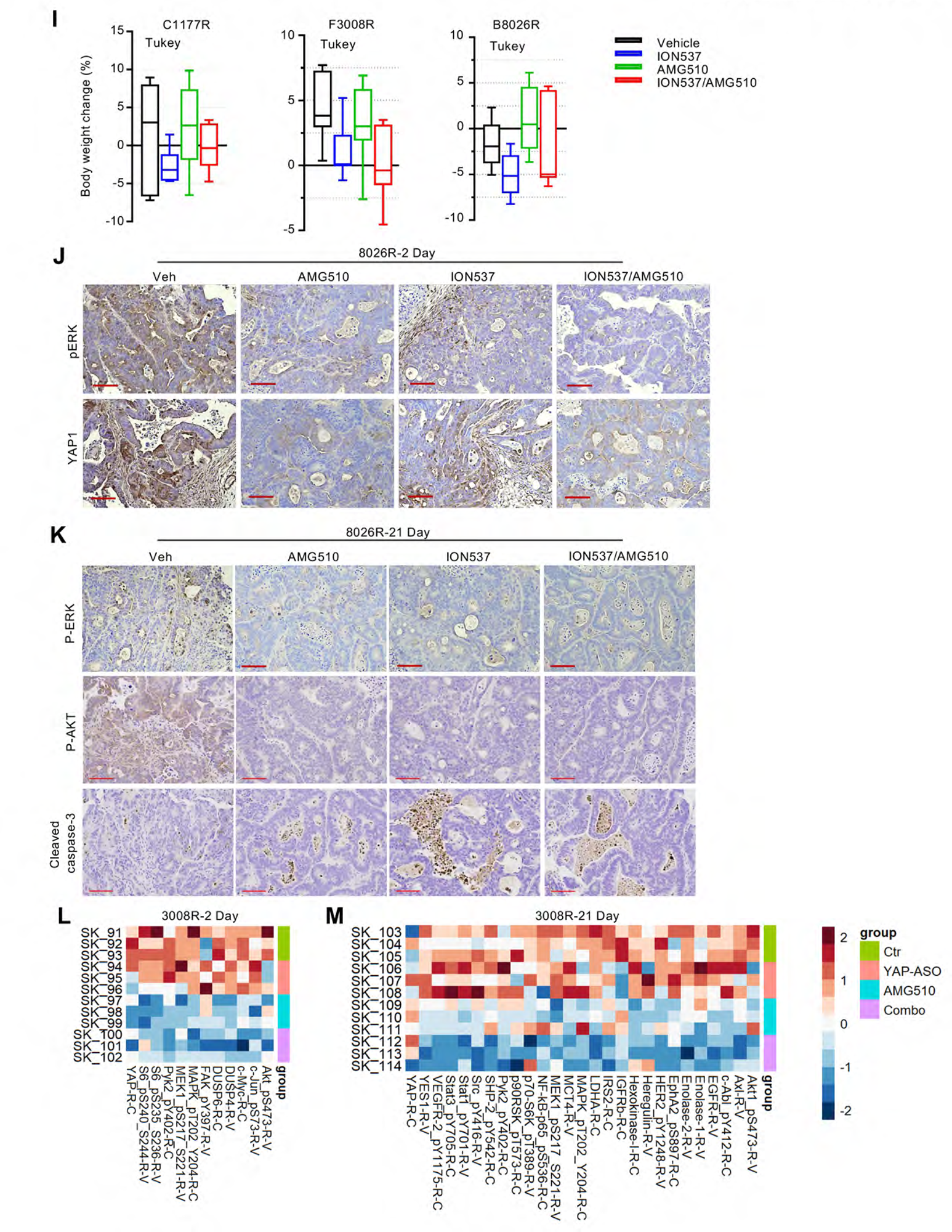

